# A three-genome ultraconserved element phylogeny of Cryptophytes

**DOI:** 10.1101/2023.09.15.557987

**Authors:** Matthew J. Greenwold, Kristiaän Merritt, Tammi L. Richardson, Jeffry L. Dudycha

## Abstract

Cryptophytes are single celled protists found in all aquatic environments. They are composed of a heterotrophic genus, *Goniomonas*, and a largely autotrophic group comprising many genera. Cryptophytes evolved through secondary endosymbiosis between a host eukaryotic heterotroph and a symbiont red alga. This merger resulted in a four-genome system that includes the nuclear and mitochondrial genomes from the host and a second nuclear genome (nucleomorph) and plastid genome inherited from the symbiont. Here, we make use of different genomes (with potentially distinct evolutionary histories) to perform a phylogenomic study of the early history of cryptophytes. Using ultraconserved elements from the host nuclear genome and symbiont nucleomorph and plastid genomes, we produce a three-genome phylogeny of 91 strains of cryptophytes. Our phylogenetic analyses find that that there are three major cryptophyte clades: Clade 1 comprises *Chroomonas* and *Hemiselmis* species, Clade 2, a taxonomically rich clade, comprises at least twelve genera, and Clade 3, comprises the heterotrophic *Goniomonas* species. Each of these major clades include both freshwater and marine species, but subclades within these clades differ in degrees of niche conservatism. Finally, we discuss priorities for taxonomic revision to Cryptophyceae based on previous studies and in light of these phylogenomic analyses.

## Introduction

Cryptophytes (= cryptomonads = Cryptophyceae) are a group of single-celled eukaryotic protists that are either heterotrophs, autotrophs, or mixotrophs (Adl et al., 2012; Hoef-Emden and Archibald, 2016). Autotrophic and mixotrophic cryptophytes evolved from secondary endosymbiosis when an unknown eukaryote host engulfed a red alga symbiont. This merger provided cryptophytes the ability to harvest energy through photosynthesis (Douglas and Penny, 1999; Douglas et al., 2001; Gould et al., 2008). Secondary endosymbiosis also resulted in cryptophytes acquiring a second nuclear genome referred to as the nucleomorph and a plastid genome forming a four-genome system (Gillott and Gibbs, 1980; Ludwig and Gibbs, 1985; Douglas et al., 2001; Curtis et al., 2012). Cryptophytes contain chlorophyll *a* and *c_2_* as well as a unique pigment-protein complex referred to as cryptophyte phycobiliproteins (Hill and Rowan, 1989; Hoef-Emden and Archibald, 2016). Cryptophyte phycobiliproteins differ among species and are classified based on the maximum absorption peak and pigment, which can be phycoerythrin (PE; appearing purple to orange; for example, blueish-red of Cr-PE566 to yellowish-red of Cr-PE545 and Cr-PE555) or phycocyanin (PC; green to blue; for example, Cr-PE545 or Cr-PC630). Nine distinct cryptophyte phycobiliproteins are known, allowing cryptophytes (in combination with chlorophylls and carotenoids) to display a fascinating breadth of colors and make use of the light in the spectrum not used by chlorophyll (Glazer and Wedemayer, 1995; Magalhaes et al., 2021).

Cryptophyta have long been recognized as a phylum consisting of two classes that include a heterotrophic group, Goniomonadea, and a mostly autotrophic group, Cryptophyceae (Hoef-Emden and Archibald, 2016). The last major revision of Cryptophyta published by Clay et al., in 1999 using available ultrastructural, biochemical, and molecular data, divided the phylum into two classes, three orders, and six families, including three newly proposed families. However, there have been several suggested additions to the number of families/genera, reductions of genera, and other minor revisions based on molecular and morphological data (Clay et al., 1999; Hoef-Emden and Melkonian, 2003; Majaneva et al., 2014; Majaneva et al., 2016; Shiratori and Ishida, 2016; Daugbjerg et al., 2018; Hoef-Emden, 2018; Altenburger et al., 2020). Recently, however, Adl et al. (2019) have “informally” suggested that cryptophytes (cryptomonads) form the rank Cryptophyceae within the Cryptista along with kathablepharids and *Palplitomonas bilix* (Yabuki et al., 2014). Adl et al. (2019) made these “informal” suggestions only if sufficient evidence that a group or clade (of eukaryotes) could be considered monophyletic. Thus, they suggested that the heterotroph class (previously Goniomonadea) and the autotroph class (previously Cryptophyceae) comprise the monophyletic clade Cryptophyceae. More recently, Yazaki et al. (2022) performed a phylogenomic analysis of unicellular eukaryotes and found that along with *Palpitomonas bilix* and *Hemiarma marina* (sister taxon to Goniomonadea; Shiratori and Ishida, 2016) that *Microheliella maris* forms a deep branch of Cryptista and together should be referred to as Pancryptista (Yabuki et al., 2014). Here, we will examine how lower taxonomic levels of Cryptophyceae may fit into the larger eukaryotic hierarchy outlined by Adl et al. (2019). First, we will review the Clay et al. (1999) classification scheme which is based on the historical term “Cryptophyta” (cryptomonads) being viewed as a phylum.

The class Goniomonadea with one order, Goniomonadida, consists solely of the heterotrophic genus *Goniomonas*. *Goniomonas* are non-photosynthetic cryptophytes that do not possess a nucleomorph or plastid genome as seen in other Cryptophyta (Kugrens and Lee, 1991; Kim and Archibald, 2013; Cenci et al., 2018). The second class of Cryptophyta is Cryptophyceae, which possess evidence of secondary endosymbiosis including a vestigial nucleus (the nucleomorph) and a plastid (Clay et al., 1999; Archibald, 2007). Thus, these Cryptophytes contain two genomes derived from the ancestral endosymbiont. The order Cryptomonadales was described by Clay et al., (1999) as consisting of two families. Cryptomonadaceae included the genus *Cryptomonas* and Campylomonadaceae included *Campylomonas* and *Chilomonas* genera. However, this order was reduced to one genus, *Cryptomonas*, because distinguishing morphological features of *Campylomonas* and *Chilomonas* were later shown to be a dimorphic state of *Cryptomonas* (see Hoef-Emden and Melkonian, 2003). Furthermore, nuclear and nucleomorph rDNA phylogenies supported the single-genus hypothesis (Hoef-Emden et al., 2002). A recent study (Daugbjerg et al., 2018), placed a third novel family, Baffinellaceae, in the order Cryptomonadales based mainly on the presence of a cryptophyte phycoerythrin 566 phycobiliprotein which had been considered a defining trait of *Cryptomonas*. Baffinellaceae has one named genus (*Baffinella*) and species; *Baffinella frigidus* (previously: *Unidentified* sp. CCMP2045 and CCMP2293). Interestingly, while these strains are sister taxa and identically named, CCMP2045 has Cr-PC566 while CCMP2293 has Cr-PE545 phycobiliproteins (Daugbjerg et al., 2018; Cunningham et al., 2019).

The third order of Cryptophyta described by Clay et al. (1999), Pyrenomonadales, is the most taxonomically rich with four families and twelve genera. The family Pyrenomonadaceae is composed of the genera *Rhodomonas*, *Rhinomonas*, and *Storeatula*. The family Geminigeraceae is composed of *Geminigera*, *Teleaulax*, *Hanusia*, *Guillardia*, and *Proteomonas*. The family Chroomonadaceae is composed of *Chroomonas*, *Falcomonas,* and *Komma*. The last Pyrenomonadales family, Hemiselmidaceae has only one genus, *Hemiselmis*. Molecular phylogenies have contradicted the monophyly of Geminigeraceae and indicate that it is composed of at least three clades (Hoef-Emden 2008; Cunningham et al., 2019). Furthermore, molecular phylogenies indicate Hemiselmidaceae may be a nested clade among one or more Chroomonadaceae clades (for example, see Hoef-Emden 2008; 2018; Cunningham et al., 2019).

Molecular data used for phylogenetic studies have grown from a handful of genes produced using Sanger sequencing to whole genome datasets (exons, introns, and conserved noncoding regions) using next-generation sequencing technologies. There are many approaches to harvesting genome-wide data for phylogenetics including transcriptome sequencing (Wang et al., 2009), whole genome sequencing (Lam et al., 2012) and reduced-representation genome sequencing (also referred to as target enrichment using molecular probes or baits; Faircloth et al., 2012). However, there are concerns with the first two methods including the difficulty of working with RNA for transcriptomics and the large monetary expense for whole genome sequencing. Target enrichment methods such as ultraconserved elements (UCEs) are a cheaper alternative to whole genome sequencing, have been shown to work on degraded DNA from historic museum specimens, and are phylogenetically informative on different timescales (Blaimer et al., 2016; Faircloth et al., 2012). Ultraconserved elements (UCEs) were first recognized in mammals and were defined as stretches of at least 200 base pairs of DNA that are highly conserved among a group of taxa (Bejerano et al., 2004). Ultraconserved elements are not necessarily protein coding regions, but can represent any region of the genome that is conserved and not repetitive (Bejerano et al., 2004). The utility of UCEs for phylogenetics is that the core region is conserved and is the target for a molecular probe or “bait” and when DNA sequencing is extended in either direction of the core region, the number of phylogenetic informative sites will increase and can be used on species separated by hundreds of millions of years (Faircloth et al., 2012). Indeed, UCEs have been used to study the evolutionary history of vertebrates, invertebrates, and have also been proposed as a method for studying phylogenetics of Alveolata (Baca et al., 2017; Branstetter et al., 2017; Crawford et al., 2012; Gilbert et al., 2015; McCormack et al., 2013; Mills et al., 2023; Parada et al., 2021; Rubanov et al., 2016; Starrett et al., 2017; Wood et al., 2020).

Here, we designed UCEs for the nuclear, nucleomorph, and plastid genomes of cryptophytes and then use them to enrich genome-wide data from 91 strains of cryptophytes and a haptophyte (*Emiliania huxleyi* CCMP373) to ascertain the evolutionary history of cryptophytes. The strains in our data set represent 15 genera and we present biochemical and habitat data for nearly all strains (Cunningham et al., 2019). We find that classical taxonomy based on morphology does not consistently provide distinguishing features that reflect the evolutionary relationships of Cryptophyceae we ascertained from genomic data.

## Methods

### Culture conditions, phycobiliprotein classes, and DNA extractions

Cryptophyte strains were either purchased from culture centers or collected in North Carolina, USA and South Carolina, USA (see Supplementary Material Table S1 for details). Cultures were grown in incubators under a 12h:12h light:dark cycle in media and at temperatures appropriate for each strain. Cryptophyte phycobiliprotein classes for 33 strains are from Cunningham et al. (2019). Phycobiliprotein classes for 46 additional strains were obtained following Cunningham et al. (2019). Culture centers, sample locations, incubation temperatures, media, and cryptophyte phycobiliprotein classes are listed in Supplementary Material Table S1.

To obtain samples for DNA extraction, algal cultures were harvested at dense log-phase and centrifuged in 500 mL bottles for 30 minutes at ∼7,000 RPM in a Beckman Coulter JA-10 fixed angle rotor. The resulting cell pellets were resuspended and transferred to 2.0 mL microcentrifuge tubes. The 2.0 mL tubes were spun for 12 minutes at ∼3,000 RPM in a microcentrifuge and, after decanting the liquid, cell pellets were stored at -80°C. DNA extraction was carried out with either Qiagen DNeasy Plant Mini Kit according to the manufacturer’s instructions or the CTAB method following Curtis et al. (2012).

### UCE probe design and sequencing

Ultraconserved element (UCE) (Supplementary Material Data S1) de novo probes for cryptophytes were designed using phyluce (release 1.5.0: March 29, 2017; Faircloth et al., 2012 and Faircloth, 2016). Probe design was conducted separately for the nuclear, nucleomorph and plastid genomes. For each genome we retained conserved loci that were found in at least two taxa. For the nuclear genome, we downloaded the *Guillardia theta* CCMP3327 and *Baffinella frigidus* CCMP2293 genomes from the Department of Energy (DOE) Joint Genome Institute (JGI) genome portal and used them as reference genomes for ultraconserved element (UCE) probe design (Curtis et al., 2012; Daugbjerg et al., 2018; Nordberg et al., 2014). We used Phyluce version 1.5.0 and *G. theta* CCMP3327 as the base genome to identify, extract and validate 7,112 nucleotide probe sequences from 1,868 conserved nuclear genome loci (Faircloth et al., 2012; Faircloth, 2016). For the nucleomorph genome, we also used *Guillardia theta* CCMP3327 (NCBI Accession #s AF165818, NC_002753, and NC_002751) as the base genome and *Chrooomonas mesostigmatica* CCMP1168 (NCBI Accession #s CP003680, CP003681, and CP003682), *Baffinella frigidus* CCMP2293, *Cryptomonas paramecium* CCAP977/2a (NCBI Accession #s NC_015329, NC_015330, and NC_015331), and *Hemiselmis andersenii* CCMP 644 (NCBI Accession # NC_009977, NC_009978, and NC_009979) genomes to produce 1,371 probes from 192 loci. *Guillardia theta* CCMP3327 (NCBI Accession # NC_000926) was used as the base genome to identify 1,436 UCE probes from 207 plastid loci in conjunction with *Baffinella frigidus* CCMP2293, *Cryptomonas paramecium* CCAP977/2a (NCBI Accession # NC_013703), *Rhodomonas salina* CCMP1319 (NCBI Accession # EF508371), and *Teleaulax amphioxeia* HACCP-CR01 (NCBI Accession # KP899713). The final probe datasets from the three genomes were combined and screened for duplicate probe sequences.

Ultraconserved element probes (Supplementary Material Data S1) designed for cryptophytes were synthesized by RAPiD Genomics (Gainesville, Florida). DNA samples were sent to RAPiD Genomics for UCE sequence capture using 150 bp paired-end Illumina sequencing. Sequence capture and Illumina sequencing were performed using the same DNA samples for all three genomes. The nuclear and nucleomorph Illumina sequencing was carried out jointly while the plastid genome was sequenced separately. Raw sequencing data is available under the NCBI BioProject # PRJNA984198.

### UCE processing

Illumina sequence data was processed using the phyluce pipeline (release 1.6.6: October 8^th^, 2019; Faircloth et al., 2012 and Faircloth, 2016). Adapter and quality trimming were performed using illumiprocessor (Faircloth, 2013; Bolger et al., 2014). DNA assembly was performed using Velvet (version 1.2.10) and ABySS with kmer values of 35-65 and Spades (Zerbino and Birney, 2007; Simpson et al., 2009; Bankevich et al., 2012). For each DNA assembly, we filtered UCEs by removing those that were found in fewer than three strains. The assembly we chose for each genome was the one with the highest number of UCE loci and nucleotides. Based on these criteria, the Spades assembly was selected for both the nuclear and nucleomorph genomes while the Velvet (kmer = 45) assembly was selected for the plastid genome. Following DNA assembly, we identified enriched UCE loci for each probe set (nuclear, nucleomorph, and plastid), removed potential paralogs and extracted enriched UCE loci using LASTZ and a minimum coverage and identity of 80% (Harris, 2007) using the phyluce “phyluce_assembly_match_contigs_to_probes” script.

### UCE Filtering

Using the selected DNA assemblies for each genome (see above), we evaluated the effect of additional filtering steps on the topology and support values using Maximum likelihood analyses (see below). We first evaluated the homogeneity of base composition of different taxa for each dataset (nuclear, nucleomorph, and plastid) and the combined dataset using a chi-square test implemented in PAUP* 4.0a (build 169; Swofford, 1998). All datasets failed the chi-square test; therefore, we evaluated if the removal of different taxa or combinations of taxa up to five strains improved the results. The removal of any single taxon or combination of taxa still resulted in a failed chi-square test. We next evaluated if certain UCE loci that had extreme GC content altered the topology or support values. For each dataset, we removed GC content outliers by removing loci that fell outside of 1.5 standard deviations of the mean GC content for each dataset. We also evaluated how the removal of loci with the bottom 5% number of parsimony informative (PI) sites, and those with a high risk of nucleotide saturation effect phylogenetic results (Duchene et al., 2022). The number of parsimony informative sites and GC content for each UCE locus was calculated using the AMAS program and risk for high nucleotide saturation for each locus was evaluated using the program PhyloMAd (Borowiec, 2016; Duchene et al., 2018; See Supplementary Material Table S2 for results). We also evaluated the effect of removing the haptophyte outgroup (*Emiliania huxleyi* CCMP373) as haptophytes do not represent the closest relative to cryptophytes (Adl et al., 2019). In summary, four filters were applied: 1) removal of loci with a low number of PI sites; 2) removal of loci with a high risk of nucleotide saturation; 3) removal of loci with extreme GC content outside 1.5 standard deviations of the mean; 4) removal of the haptophyte outgroup. All filters were applied in a pairwise fashion to evaluate the best combination of filtering steps for this UCE dataset. While the topology did not differ among most filtered datasets, the bootstrap support values did differ among datasets. However, the filtering of loci with extreme GC content and a low number of PI sites resulted in *Urgorri complanatus* (BEA0603) becoming unstable (see Results). Overall, the inclusion of the haptophyte outgroup and removal of loci with a low number of PI sites resulted in the strongest bootstrap support values and was therefore used for all subsequent phylogenetic analyses.

### Phylogenetics

Phylogenetic analyses were performed using both maximum likelihood (ML) with a concatenated alignment, and the multi-species coalescent (MSC)-based approach. The MSC uses single-locus (gene) trees of each UCE locus to produce a combined species tree. We performed maximum likelihood analyses using RAxML v. 8.0.19 and MSC using ASTRAL-III (Stamatakis, 2014; Zang et al., 2018). We used UCEs from all three genomes (nuclear, nucleomorph, and plastid) and maximum likelihood to produce a total evidence nucleotide tree (TENT; Figure 1; Supplementary Material Data S2 for a tree nexus file). We also constructed a nuclear (Supplementary Material Fig. S1), nucleomorph (Supplementary Material Fig. S2), and plastid (Supplementary Material Fig. S3) individual maximum likelihood phylogenies.

**Figure 1.**
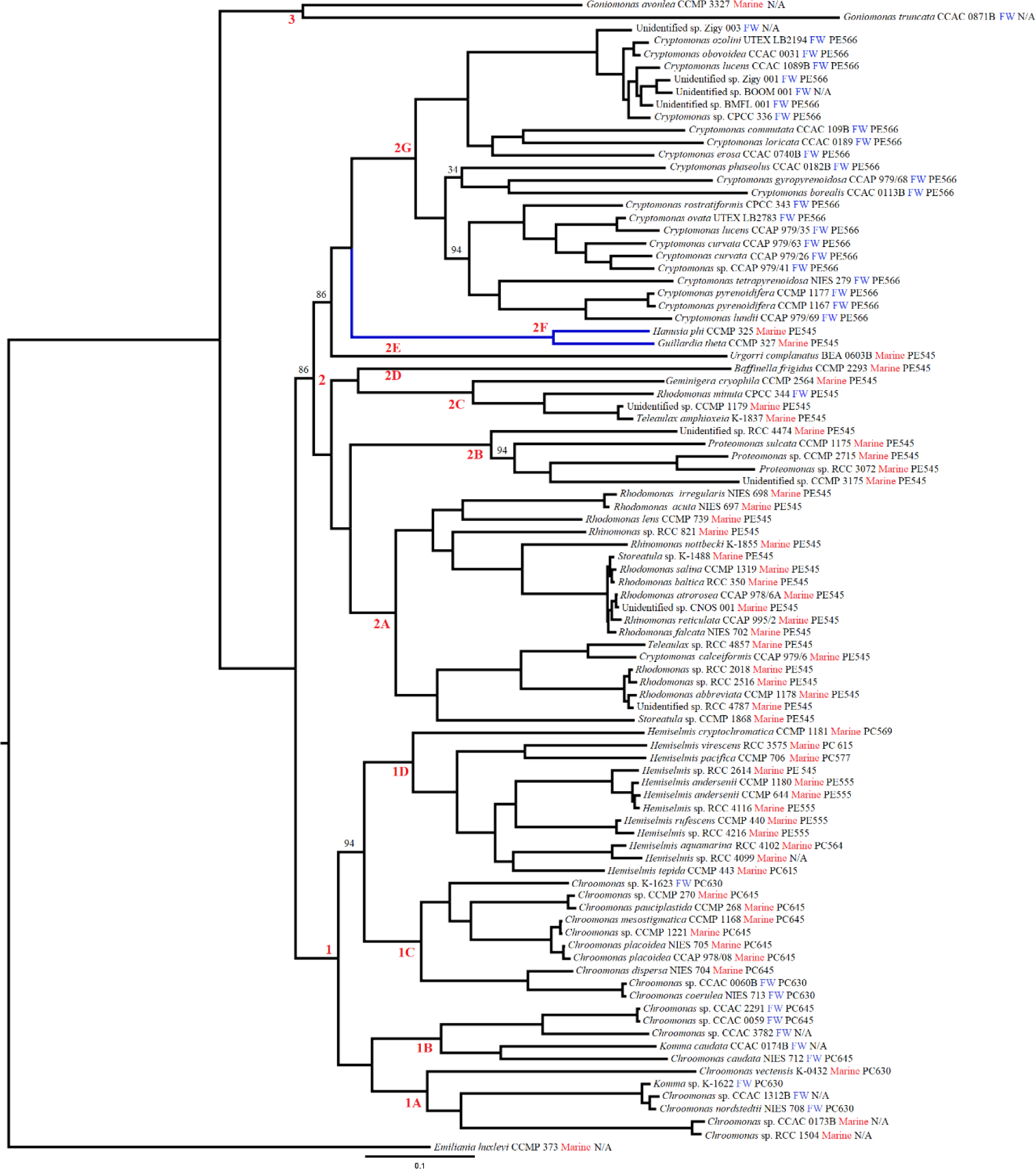
Maximum likelihood phylogeny. Bootstrap support values are listed for branches with less than 100% bootstrap support. Clade designations are listed for major branches in the phylogeny. Species names as well as the habitat and cryptophyte phycobiliprotein type are listed for each taxon (see also Supplementary Material Table S1). The highlighted branch in blue represents discordance between maximum likelihood and multispecies coalescent methods.

MAFFT v7.471 was used to produce an alignment that was trimmed using the default settings for Gblocks (Castresana, 2000; Katoh et al., 2002; Katoh and Standley, 2013). ModelTest-NG v0.2.0 (Flouri et al., 2014; Darriba et al., 2020) was used to select the best-fit DNA substitution model of GTR+I+G4 based on AIC, BIC, and AICc for all four datasets. A maximum likelihood phylogeny was constructed using RAxML v. 8.0.19. We used the autoMRE method for bootstrap replicates which were then mapped on to the best scoring phylogeny produced from 200 thorough maximum likelihood replicates (Stamatakis, 2014).

Multi-species coalescent (MSC)-based phylogenomic analyses were conducted using ASTRAL-III v.5.7.7 (Zang et al., 2018). Single-locus trees of each UCE locus were constructed using RAxML v. 8.0.19 and the -f a option with 200 replicates (Stamatakis, 2014). RAxML best-scoring maximum likelihood single-locus trees and bootstrap replicates were provided to ASTRAL-III. We performed the ASTRAL-III analyses after removing low-support branches (below 10% bootstrap support) as specified in the manual (Zang et al., 2018; https://github.com/smirarab/ASTRAL/blob/master/astral-tutorial.md). ASTRAL-III local posterior probabilities are shown in Supplementary Material Fig. S4.

Species tree scoring was performed using the -q option, which is the fraction of quartet trees that are in the species tree. Quartet scoring was performed using single-locus trees based on the evolutionary history of cryptophytes (nuclear single-locus trees and nucleomorph + plastid single-locus trees) to evaluate the contribution of phylogenetic discordance from different genomes (with distinct evolutionary history) to the maximum likelihood and MSC phylogenies. Furthermore, we used DiscoVista and the relative frequency analysis (Sayyari et al., 2018) to analyze the quartet support for alternative branching (discordance) between our phylogenies. Relative frequency analysis was performed separately for the nuclear single-locus trees and the single-locus trees from the nucleomorph and plastid genomes. To calculate quartet scores for the single-locus trees of the combined gene-tree dataset of the nucleomorph + plastid genomes, we removed *Goniomonas* strains from the maximum likelihood and MSC phylogenies as *Goniomonas* strains lack a nucleomorph and plastid genome (Cenci et al., 2018). The *Goniomonas* tips were removed using the R package ape (Paradis and Schliep, 2019).

A polytomy test using quartet frequencies for the subclades of Clade 2 (2A, 2B, 2C, 2D, 2E, 2F, 2G) was performed with the ASTRAL package (Sayyari and Mirarab, 2018). The polytomy test assesses the null hypothesis that “the estimated single-locus tree quartets around the branch β support all three NNI rearrangements around the branch in equal numbers” or, simply, if the clade is better represented as a polytomy (Sayyari and Mirarab, 2018). We only considered the major branches of the maximum likelihood and MSC phylogeny as shown in Figure 2A and 2C.

**Figure 2.**
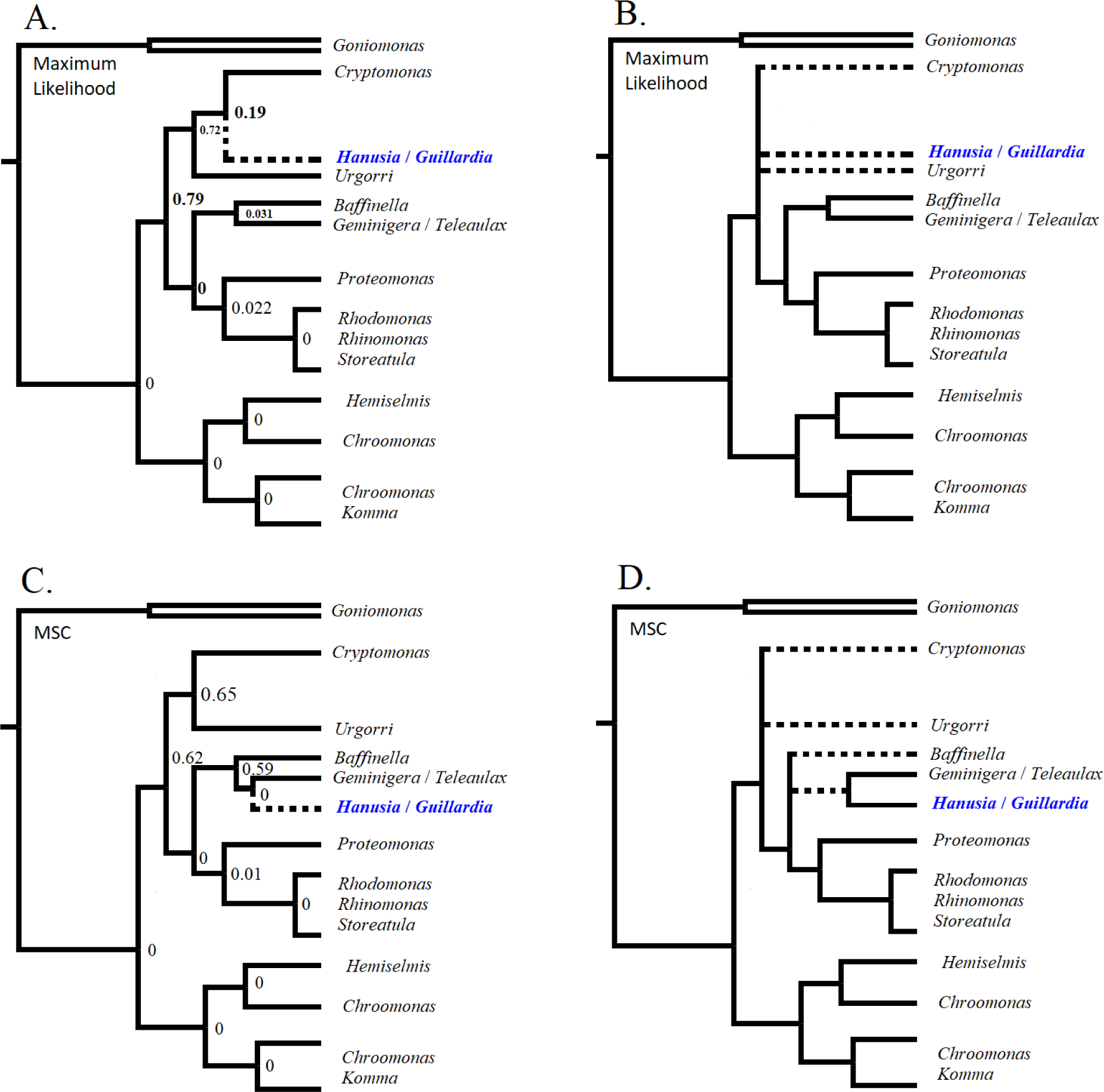
Simplified versions of maximum likelihood and multispecies coalescent (MSC) phylogenies. Dotted lines represent uncertain branching based on phylogenetic discordance between methods or polytomy test results (Sayyari and Mirarab, 2018). A; Simplified maximum likelihood phylogeny (Figure 1) with polytomy test values for each branch; B; Simplified maximum likelihood phylogeny based on polytomy test results; C; Simplified multispecies coalescent (MSC) phylogeny (Supplementary Material Fig. S4) with polytomy test values for each branch; D; Simplified multispecies coalescent (MSC) phylogeny based on polytomy test results.

We evaluated the possible influence of long branch attraction (LBA) on the discordance observed between the maximum likelihood and MSC phylogenies using TreeShrink 1.3.9 (Mai and Mirarab, 2018). We evaluated the single TENT maximum likelihood phylogeny (Figure 1) and the set of single-locus trees used for MSC analyses (Supplementary Material Fig. S4) using default settings. For the TENT phylogeny, per-gene analysis was performed and per-species was used for the single-locus trees.

## Results

Using next-generation sequencing, we sequenced 1,631 nuclear loci, 125 nucleomorph loci, and 187 plastid loci from 89 strains of plastid-bearing cryptophytes. These data were combined with UCE nuclear data from two heterotrophic *Goniomonas* strains (*Goniomonas* lacks a nucleomorph and plastid), and UCE nuclear and plastid data for one haptophyte species (*Emiliania huxleyi* CCMP373; which lacks a nucleomorph) to produce a total evidence data set consisting of 1,943 loci with an alignment length of 349,254 characters, 173,842 parsimony-informative sites, and an average of 89 parsimony-informative sites per locus (Table 1).

**Table 1.**
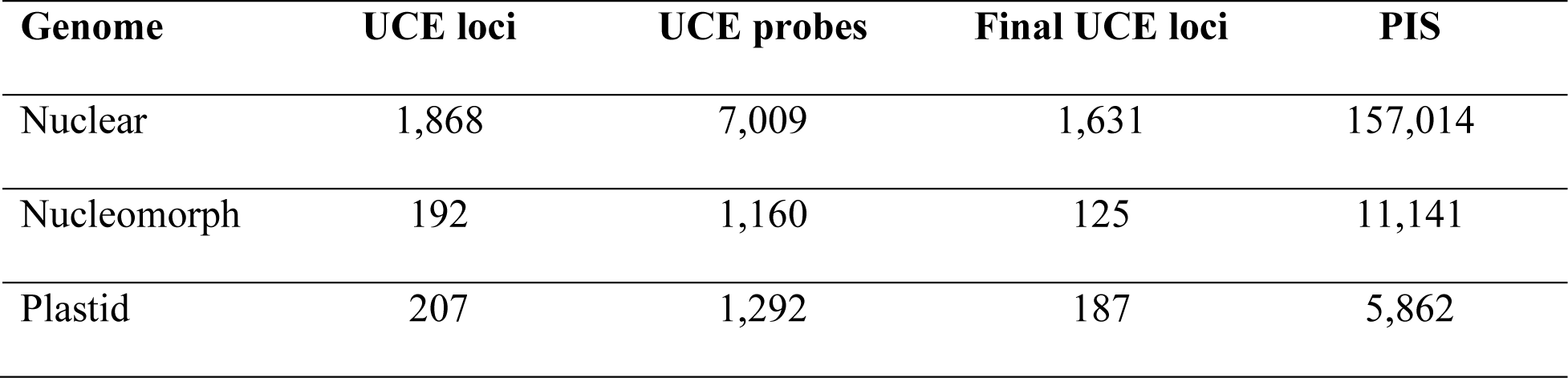
Number of UCE loci, UCE probes, amplified UCE loci, and Parsimony Informative Sites (PIS) for each genome.

| Genome | UCE loci | UCE probes | Final UCE loci | PIS |
| --- | --- | --- | --- | --- |
| Nuclear | 1,868 | 7,009 | 1,631 | 157,014 |
| Nucleomorph | 192 | 1,160 | 125 | 11,141 |
| Plastid | 207 | 1,292 | 187 | 5,862 |

The maximum likelihood phylogeny (Figure 1) supports the sister relationship of photosynthetic cryptophytes to heterotrophic *Goniomonas* strains (Goniomanadea; Clade 3). The photosynthetic cryptophytes (Cryptophyceae) form two main clades: Clade 1, consisting of *Chroomonas* (*Komma*) and *Hemiselmis* and a second, more genus-rich clade (Clade 2), with eleven genera including the speciose *Cryptomonas* and *Rhodomonas*. Within Clade 1, we find one *Hemiselmis* subclade and three *Chroomonas* subclades (1A-1D). Within Clade 2, we find that there are *Rhodomonas* (plus *Storeatula* and *Rhinomonas,* 2A), *Proteomonas* (2B), *Geminigera*/*Teleaulax*/*Rhodomonas* (2C), *Hanusia*/*Guillardia* (2E), and *Cryptomonas* (2G) subclades and two monospecific subclades comprised of *Urgorri complanatus* (BEA0603) and *Baffinella frigidus* (CCMP2293). *Urgorri complanatus* (BEA0603) may be an unstable branch as one of our UCE filtering evaluations (removal of loci with extreme GC content and a low number of PI sites) resulted in *U. complanatus* forming a monospecific outgroup to the entire Clade 2, albeit with low bootstrap support.

There are several apparent taxonomic discrepancies within Clade 2. We find that one named *Cryptomonas* species (*C*. *calceiformis* CCAP979/6) is found within subclade 2A of the Pyrenomonadaceae family. This “*Cryptomonas*” strain is marine and possesses the PE545 phycobiliprotein type seen in other Pyrenomonadaceae family members. All other *Cryptomonas* in this study were isolated from freshwater and possess the Cr-PE566 PBP (Supplementary Material Table S1). Subclade 2A also contains a strain identified as *Teleaulax* whereas the other strain identified as *Teleaulax* is in subclade 2B. However, subclade 2B appears to be an unusual group of genera consisting of strains identified as *Rhodomonas*, *Geminigera*, and *Teleaulax*. All these strains possess the Cr-PE545 PBP. In fact, except for subclade 2G, all strains in Clade 2 contain the Cr-PE545 PBP (Figure 1; Supplementary Material Table S1).

The maximum likelihood phylogeny (Figure 1) presented here does not support the monophyly of the family Geminigeraceae (*Geminigera*, *Teleaulax*, *Hanusia*, *Guillardia*, and *Proteomonas*) based on morphology and PBP type (Clay et al., 1999). Strains from these genera are found in at least three phylogenetic subclades. We find that subclade 2F (*Guillardia*/*Hanusia*) is grouped with the family Cryptomonadaceae (Cryptomonas; subclade 2G), while *Proteomonas* strains (subclade 2B), a *Geminigera* strain (subclade 2C), and *Teleaulax* strains (subclades 2A and 2C) are more closely related to the Pyrenomonadaceae family (*Rhodomonas*, *Rhinomonas*, and *Storeatula*; subclade 2A). We also do not find support for the monophyly of the Chroomonadaceae family consisting of *Chroomonas* and *Komma* strains (subclades 1A-1C) to the exclusion of the Hemiselmidaceae family (subclade 1D; *Hemiselmis*).

We also constructed a multispecies coalescent (MSC)-based phylogeny using single-locus trees produced from the total evidence data set of 1,943 UCE loci (Figure 2C; Supplementary Material Fig. S4). Overall, we found good agreement between the maximum likelihood phylogeny based on the concatenated alignment and MSC phylogeny. However, one major discrepancy between the two methods was seen with the placement of the clade consisting of *Hanusia* and *Guillardia* strains (Figure 1; clade 2F). The maximum likelihood phylogeny indicates that *Hanusia* and *Guillardia* group with *Cryptomonas* strains, while the MSC approach indicates that they are more closely related to *Baffinella*, *Proteomonas*, *Geminigera*, *Teleaulax,* and *Rhodomonas* strains (Figure 2). Physiologically, the latter makes sense as *Hanusia* and *Guillardia* both possess the PE545 cryptophyte phycobilin and inhabit marine environments as do *Rhodomonas* strains. Furthermore, this generally agrees with the morphological based classification of Clay et al. (1999), where the Family Geminigeraceae includes five genera (*Geminigera*, *Teleaulax*, *Hanusia*, *Guillardia*, and *Proteomonas*).

To further analyze the discordance between the maximum likelihood and MSC phylogeny, we calculated the ASTRAL-III normalized quartet scores (fraction of quartet trees that support a phylogeny; Zhang et al., 2018). The maximum likelihood phylogeny (0.7736) has a slightly lower quartet score than the MSC phylogeny (0.7772) based on single-locus trees from all three genomes (nuclear, nucleomorph, and plastid). This held true even when we separated the nuclear single-locus trees and nucleomorph + plastid single-locus trees (Table 2). The strongest quartet support for a phylogeny are the nuclear single-locus trees for the MSC phylogeny (Table 2).

**Table 2.**
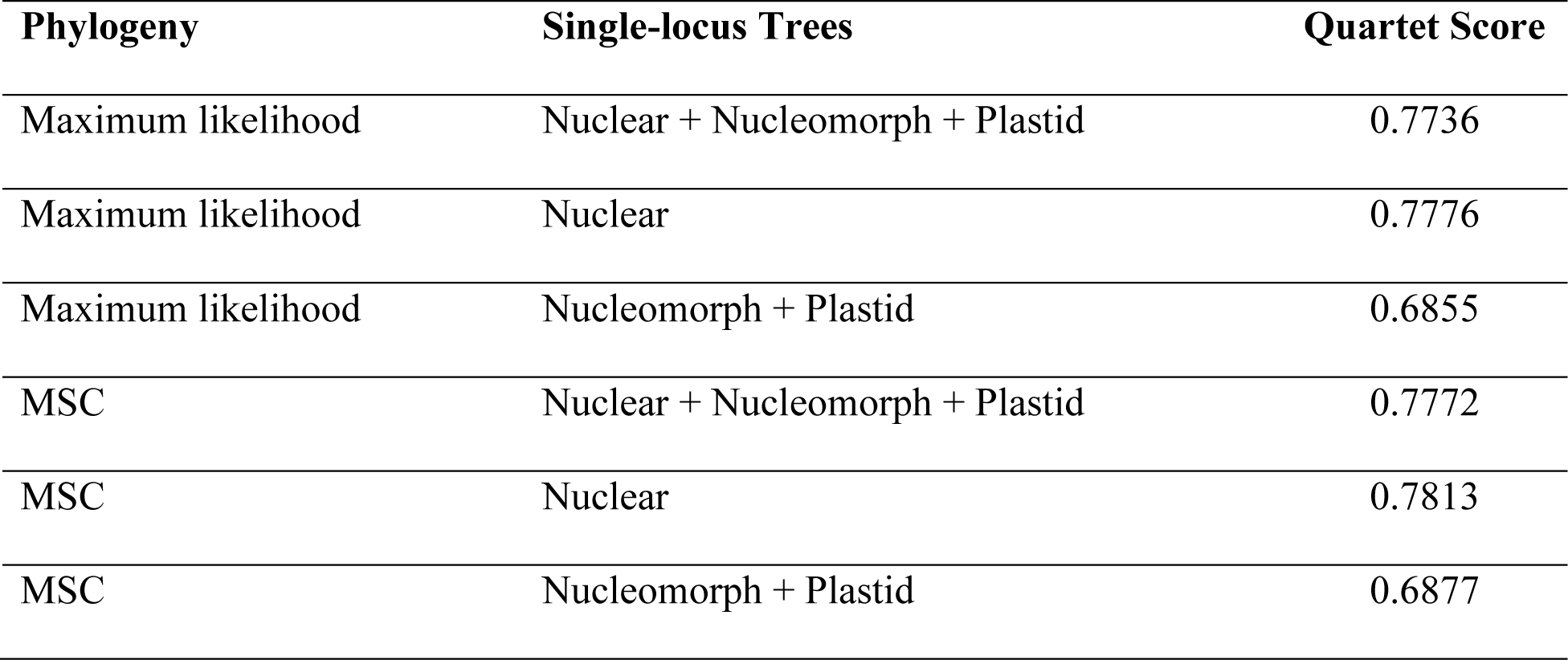
Quartet scores for maximum likelihood and MSC phylogenies using different single-locus trees datasets.

| Phylogeny | Single-locus Trees | Quartet Score |
| --- | --- | --- |
| Maximum likelihood | Nuclear + Nucleomorph + Plastid | 0.7736 |
| Maximum likelihood | Nuclear | 0.7776 |
| Maximum likelihood | Nucleomorph + Plastid | 0.6855 |
| MSC | Nuclear + Nucleomorph + Plastid | 0.7772 |
| MSC | Nuclear | 0.7813 |
| MSC | Nucleomorph + Plastid | 0.6877 |

We also used DiscoVista (Sayyari et al., 2018) to study the discordance between single-locus trees and our maximum likelihood (Figure 3) and MSC phylogenies (Supplementary Material Fig. S5) using a relative frequency analysis. We separated our single-locus trees into two sets: 1) nuclear; and 2) nucleomorph + plastid single-locus trees based on the evolutionary history of cryptophytes. We hypothesized that the evolutionary history (secondary endosymbiosis) of cryptophytes plays a significant role in the discordance between maximum likelihood and MSC phylogenies. As branch 8 in Figure 3 indicates, there is slightly better support for the maximum likelihood phylogeny (*Hanusia*/*Guillardia* closely related to *Cryptomonas*) from the nuclear single-locus trees (10,11|7,9; ∼0.35 vs ∼0.3), while the MSC phylogeny (*Hanusia*/*Guillardia* closely related to *Geminigera/Teleaulax*; see Supplementary Material Fig. S5, branch 3) is better supported by the nucleomorph and plastid single-locus trees (10,9|4,6; ∼0.5 vs ∼0.45). Interestingly, the nucleomorph + plastid single-locus trees support an alternative topology from our maximum likelihood and MSC phylogenies where the clade comprising *Chroomonas/Hemiselmis* forms the outgroup of autotrophic cryptophytes as seen for branch 6 of Figure 3 (see also branch 12 of Supplementary Material Fig. S5).

**Figure 3.**
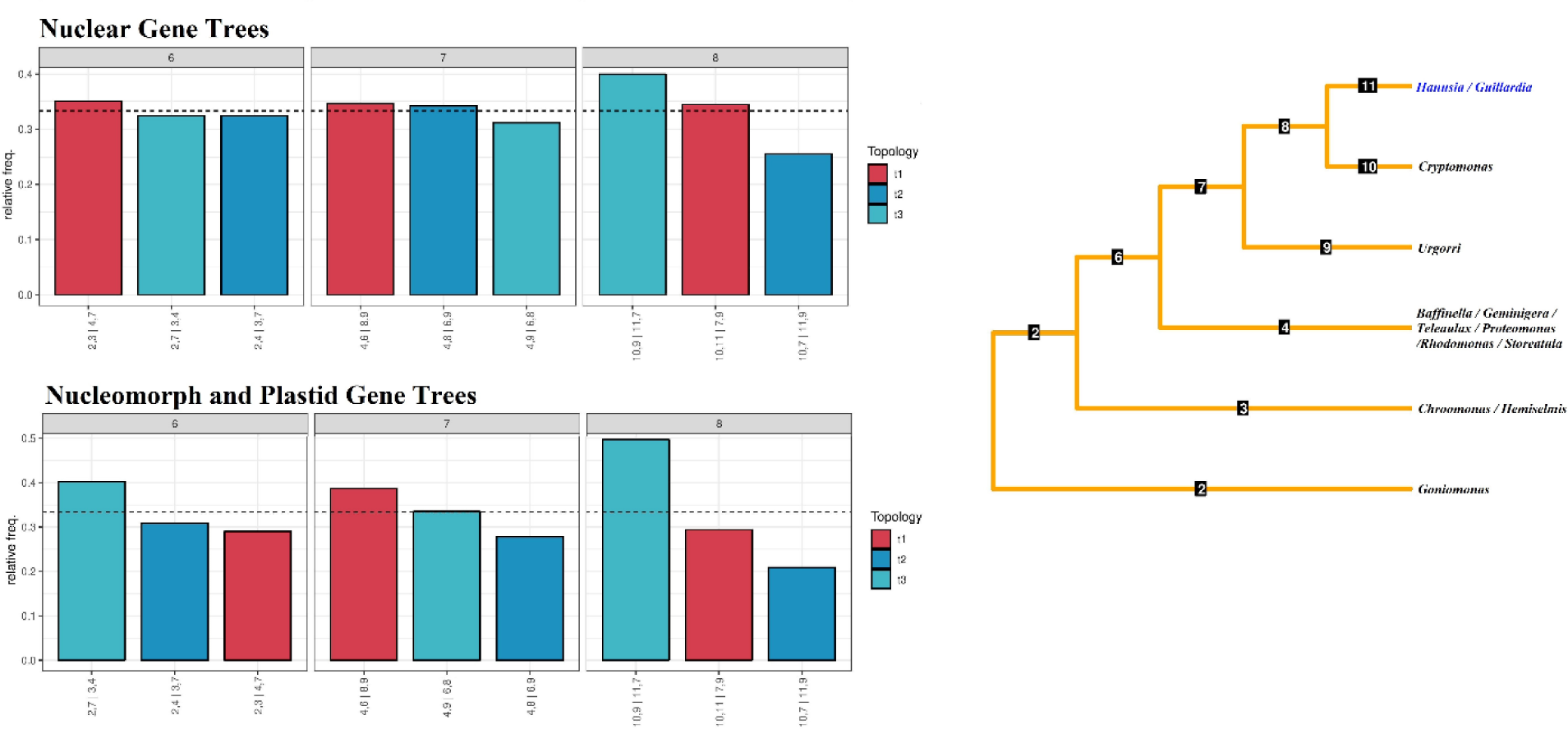
DiscoVista relative frequency analysis (Sayyari et al., 2018). The phylogeny on the right side is the DiscoVista simplified maximum likelihood phylogeny. For each internal branch of the phylogeny a relative frequency of single-locus trees that support that branch are reported for each single-locus tree dataset (nuclear and nucleomorph + plastid) based on the evolutionary history of cryptophytes. Each internal branch has four connected branches that form three different topologies. The red (t1) bars indicate relative frequency of single-locus trees that support the maximum likelihood phylogeny, and the blue (t2 and t3) bars are alternative topologies. The dotted line indicates one-third of the total number of single-locus trees. Each topology is detailed on the x-axis where the branches separated by a comma are joined.

The phylogenetic discordance seen with the placement of the *Hanusia/Guillardia* clade between the maximum likelihood and MSC phylogeny led us to evaluate if our phylogenomic data better supports a polytomy for Clade 2 (Figure 2). We performed a polytomy test using quartet frequencies and ASTRAL-III (Sayyari and Mirarab, 2018; (Zang et al., 2018). We cannot reject the null hypothesis at a 95% confidence level that Clade 2 is a polytomy as indicated by the test values listed on the simplified phylogeny in Figure 2A. Figure 2C illustrates that despite strong single-locus tree support for the MSC phylogeny, the quartets cannot reject a polytomy for *Cryptomonas*/*Urgorri* and the remaining Clade 2 subclades. This is not surprising as our UCE filtering and maximum likelihood analyses found that *Urgorri complanatus* (BEA0603) may be an unstable taxon (see Methods). Furthermore, *Baffinella*, *Geminigera*/*Teleaulax*/*Hanusia*/*Guillardia* form a polytomy with the clade of *Proteomonas* and *Rhodomonas*/*Rhinomonas*/*Storeatula*. Figures 2B and 2D illustrate the results of this test indicating that Clade 2 may not be strictly bifurcating based on this phylogenomic data and instead forms a polytomy.

Finally, we evaluated whether phylogenetic discordance may be associated with long branch attraction (LBA). Specifically, we posited that discordance between phylogenies is associated with long-branch taxa around the discordance; *Urgorri complanatus* (BEA 0603B) and *Hanusia*/*Guillardia*. The TENT maximum likelihood phylogeny and the single-locus MSC trees were evaluated for long branch taxa. For the TENT maximum likelihood concatenated phylogeny, only *Goniomonas truncata* (CCAC 0871B) was identified as a long-branch taxon. For the 1,943 single-locus trees, we found that *Hanusia phi* (CCMP325) was only found as a long-branch taxon in 2.2% of the trees, *Guillardia theta* (CCMP 327) in only 2.4% of the trees, and *Urgorri complanatus* (BEA 0603) in less than 1% of trees indicating that the taxa associated with the phylogenetic discordance are likely not significant long-branch taxa leading to the discordance.

## Discussion

### Niche Conservatism

Using phylogenomic data from the nuclear, nucleomorph, and plastid genomes of cryptophytes we constructed species-level estimates of the relationships within Cryptophyceae. While there is discordance between our species phylogenies produced using a concatenated alignment based maximum likelihood analysis and multispecies coalescent (MSC) approach, there are several evolutionary and ecological inferences that can be made based on agreement between the phylogenies. One is that there is strong niche conservatism associated with habitat (marine vs. freshwater) and phycobiliprotein type in Clade 2. Clade 2G (Figure 1) is comprised solely of freshwater cryptophytes that contain Cr-PE566 phycobiliproteins, while Clades 2A, 2B, 2D, 2E and 2F are comprised entirely of marine, Cr-PE545 cryptophytes. The only instance of within-subclade variation is a freshwater species in Clade 2C. In contrast, Clade 1 lacks this strong level of niche conservatism and is composed of both marine and freshwater species and at least 8 phycobiliprotein types representing multiple transitions of habitat and physiology including a *Hemiselmis* sp. (RCC2614) with Cr-PE545. Clades 1A and 1C suggest that *Chroomonas* species may be able to switch habitats relatively easily from marine to freshwater and freshwater to marine. Indeed, a marine and freshwater strain from Clade 1A is able to tolerate freshwater, brackish, and marine salinities while at least five marine strains from Clade 1C can also tolerate freshwater and brackish salinities (see Hoef-Emden, 2014). Furthermore, and more dynamically, Clade 1D demonstrates that *Hemiselmis* species are able to reverse phycobiliprotein types evolutionarily as noted by previous studies (Hoef-Emden, 2008; Hoef-Emden and Archibald, 2016; Greenwold et al., 2019). Both the maximum likelihood and MSC phylogenies support a single transition to freshwater by *Cryptomonas* species (Clade 2G) from an ancestral marine cryptophyte despite the results of the polytomy tests (Figure 1 and 2).

### Phylogenetic Discordance

This is not the first study that has found a close phylogenetic relationship of *Guillardia*/*Hanusia*/*Urgorri* and *Cryptomonas*. Johnson et al. (2016) found exactly this same relationship using partial rbcL sequences. Greenwold et al. (2019) also found that *Guillardia* forms a clade with *Cryptomonas* in two separate phylogenies using nuclear LSU and SSU sequences in one and plastid rbcL sequences in the other. However, our MSC phylogeny and relative frequency analysis using single-locus trees (UCE loci) from the nuclear and nucleomorph + plastid genomes do not support the close relationship of *Guillardia*/*Hanusia* and *Cryptomonas* (branch 8 Figure 3; branch 3 Supplementary Material Fig. S5) found in our maximum likelihood phylogeny (Figure 1 and Figure 2A). Why is a clade comprising *Guillardia*/*Hanusia*/*Urgorri*/*Cryptomonas* found so often and strongly supported by our maximum likelihood phylogeny, but not the MSC analysis using the same set of UCE loci (Supplementary Material Fig. S4)? One explanation may relate to the phylogenetic informativeness of some loci. The MSC phylogeny method does not take into account how many informative sites a locus contains and only considers the branching pattern of the single-locus tree. If a disproportionately high number of phylogenetically informative sites are in loci that support the *Guillardia*/*Hanusia*/*Cryptomonas* clade, then that clade would be favored in the concatenated alignment/maximum likelihood phylogeny. Finally, we can likely rule out long branch attraction (LBA) as a significant cause based on our analyses.

Reconciling the backbone of the cryptophyte phylogeny has been a persistent problem (for example, Hoef-Emden, 2008; Hoef-Emden and Archibald, 2016). Here we sought to reconcile the phylogeny backbone of the Cryptophyceae using phylogenomic data from three genomes. While we have discordance between phylogenetic methods, we can still form hypotheses about the early history of autotrophic cryptophytes. One intriguing hypothesis is serial endosymbiosis, similar to what is proposed to explain the early evolutionary history of chromist algae (Stiller et al., 2014). That hypothesis suggests that secondary, tertiary, and quaternary endosymbiosis have occurred in lineages containing plastids derived from red algae, where cryptophytes received their plastid from a red alga, then in turn provided it to ochrophytes, who then provided it to haptophytes (Stiller et al., 2014). One of the key aspects of the serial endosymbiosis hypothesis is that different lineages of heterotrophs acquired photosynthesis in this manner. Indeed, endosymbioses are likely very common (and frequent) among protists and use many modes of integration (Nowack and Melkonian, 2010). The difficulty of resolving the cryptophyte backbone could be similar in that multiple heterotroph lineages acquired (or shared) the same endosymbiont at or around the same time and that the core nuclear genome of these heterotrophs was similar enough to produce the discordance we see between our phylogenies. Indeed, red (cryptophytes) and green (chlorarachniophyte) alga-derived nucleomorph genomes are remarkably similar in that the genomes are much reduced and composed of only three chromosomes suggesting that similar evolutionary processes could have occurred for an endosymbiont shared between different heterotrophs (Douglas et al., 2001; Gilson et al., 2006). Furthermore, hybridization or cytoplasm transfer could have exacerbated the placement of *Guillardia*/*Hanusia* where the host heterotroph (nuclear genome) of *Guillardia*/*Hanusia* and *Cryptomonas* are the same, but the cytoplasm (nucleomorph and plastid genomes) was shared with *Geminigera*/*Teleaulax*. This may explain the relatively strong nuclear support of *Guillardia*/*Hanusia*/*Cryptomonas* clade and the very strong nucleomorph + plastid support of a *Guillardia*/Hanusia/*Geminigera*/*Teleaulax* clade. Another possibility is that an early *Cryptomonas* strain engulfed or merged with a common ancestor of *Geminigera*/*Teleaulax* resulting in the reticulate origin of *Hanusia*/*Guillardia*. This latter possibility is enticing if we consider that *Cryptomonas* strains can be heterotrophic (secondary loss of photosynthesis) and mixotrophic, and generally have a relatively greater cell volume than other cryptophytes (Hoef-Emden, 2005; Cunningham et al., 2019).

### Missing Taxa

Our study is limited to taxa we could obtain from culture collections and local isolation, and thus we were unable to include all described Cryptophyte genera (Supplementary Material Table S3). Existing algal culture collections emphasize marine species and isolations from the northern hemisphere, limiting our ability to include freshwater taxa, especially those from South America, Africa, and Australasia. However, most of the described genera of Cryptophyta excluded from our phylogeny have an uncertain taxonomic status and very limited published information. In addition to the two genera shown to be alternate morphs of *Cryptomonas* (*Chilomonas*, *Campylomonas*; Hoef-Emden and Melkonian, 2003), *Plagioselmis* is now considered an alternate morph of *Teleaulax* (Altenburger et al., 2020). *Isoselmis* has been proposed as a “nomen dubium,” equivalent to *Plagioselmis* (Novarino et al., 1994), and thus may also be an alternate morph of *Teleaulax* or a related genus. Three genera (*Olivamonas*, *Pseudocryptomonas*, and *Chrysidella*) no longer have any taxonomically valid species (Guiry and Guiry, 2022), and thus should be considered retired. As of June 2022, twelve genera have zero records in WebofScience.com and two or fewer records in AlgaeBase.org, suggesting that they have been seen rarely, if at all, after they were described (Guiry and Guiry, 2022). Three other genera retrieve only one or two records in WebofScience. With the exceptions of *Protocryptomonas* and possibly *Kisselevia*, these 15 rarely studied genera contain only one or two described species. Nearly all of these genera are freshwater.

Several genera have had their holotype species reclassified into other genera, leaving the generic status of remaining species unknown. *Cryptochrysis commutata* has been emended to *Cryptomonas commutata* (Hoef-Emden, 2007) and appears in Clade 2G of our phylogeny (Figure 1). *Cyanomonas americana* has been transferred to *Chroomonas* (Hill, 1991). *Cryptella cyanophora* has been renamed *Naisa cyanophora* (Molinari-Novoa et al., 2021), but no other information on this genus has been published. In addition, the heterotroph *Cyathomonas truncata* has been synonymized with *Goniomonas* (Larsen and Patterson, 1990). While not the holotype, the absence of published information on other *Cyathomonas* species calls this genus into question.

The marine genus *Hillea* has received some attention, but the only culture collection that reports having a strain did not respond to repeated inquiries. Butcher (1952) placed *Hillea* in the in the family Cryptomonadaceae suggesting that *Hillea* may group or form a sister group with *Cryptomonas* strains (Clade 2G); later Butcher (1967) classified *Hillea* as belonging to the mongeneric family Hilleaceae. Indeed, Adl et al. (2019) listed *Hillea* as *incertae sedis*. *Falcomonas daucoides*, the only species in its genus and formerly a part of *Hillea*, has been included in previous single-gene phylogenies (Clay et al., 1999; Deane et al., 2002), but we were unable to obtain a specimen for our study as it does not appear to be in any culture collections. The prior phylogenies suggest *Falcomonas* would be the sister taxon to our Clade 1 (Figure 1). Finally, *Hemiarma marina* is a recently described heterotroph most closely related to unknown environmental samples based on a ribosomal RNA gene (Shiratori & Ishida, 2016). Nishimura et al. (2020) place as a sister taxon to *Goniomonas* and outside the autotrophic cryptophytes based on a mitochondrial genome sequence.

### Taxonomic Recommendations

The taxonomic classification of cryptophytes has not been analyzed using more than a handful of genes or genes from multiple genomes. Here, we built species phylogenies using over 349,000 characters from three genomes of 91 cryptophyte strains to reconstruct the evolutionary history of cryptophytes. This allows us to reconsider the last major taxonomic revision by Clay et al. (1999) and form a proposal to revise Class Cryptophyceae considering the major taxonomic revisions of eukaryotes by Adl et al. (2019). Even with uncertainty from polytomies, our three-genome phylogeny shows that substantial taxonomic revision above the species level is necessary to reconcile taxonomy with evolutionary history. Clay et al. (1999) recognized two Cryptophyceae orders which must be reclassified as Families. Based on our phylogenies (Figure 1; Supplementary Material Fig. S4), these two families would represent Clade 1 and 2. However, the taxa contained in the Clay et al. (1999) two orders largely in conflict with our results. Clade 1 (Figure 1) could be recognized as a distinct Family named Chroomonadaceae consisting of *Hemiselmis*, *Chroomonas*, and *Komma* strains with four subfamilies consisting of three *Chroomonas* subfamilies and one *Hemiselmis* subfamily.

Within Clade 1, *Hemiselmis* (1D) is a reliable genus, unlike *Chroomonas*. The paraphyletic nature of *Chroomonas* has long been known and is not congruent with morphological data (electron microscopy) and/or biochemical data (see Hoef-Emden, 2018). Specifically, Hoef-Emden (2018) found that the *Chroomonas* subclades group based on periplast surface geometry using nuclear and nucleomorph rDNA sequences. However, contrary to Hoef-Emden (2018), we find that clade 1A and 1B have hexagonal and rectangular morphology, respectively, while clade 1C has rectangular periplast morphology. Clades 1A and 1B contain the holotypes for *Chroomonas* Hansgirg (1885; *C. nordstedtii*) and *Komma* Hill (1991; *K. caudata*) respectively and should therefore align with those genera. Each of those clades contain a single phycobiliprotein type (Cr-PC630 in 1A and Cr-PC645 in 1B). Clade 1C presents a greater challenge. All of the strains are identified as *Chroomonas*, but if that generic name is matched to Clade 1A, a new name will be needed for Clade 1C. An argument could be made for further splitting Clade 1C, since it includes both marine and freshwater taxa. However, within that subclade neither marine nor freshwater are monophyletic, and it may therefore be best to erect a single dual-habitat genus.

Clade 2 of our phylogeny (Figure 1) includes genera from the Orders Cryptomonadales and Pyrenomonadales (sans *Chroomonas*, *Komma*, and *Hemiselmis*) from Clay et al. (1999). We suggest that the order Cryptomonadales be reclassified as Cryptomonadaceae and expanded to include the genera of the order Pyrenomonadales. The order Pyrenomonadales and family Pyrenomonadaceae should be dissolved since the genus *Pyrenomonas* is now considered synonymous with *Rhodomonas* and is not often used in describing new strains (Erata and Chihara, 1989). Additionally, and as suggested by Hoef-Emden and Melkonian (2003), the Family Campylomonadaceae should be dissolved as *Campylomonas* is an alternate morphology of *Cryptomonas*. Our phylogenies also suggest that the Family Baffinellaceae (Daugbjerg et al., 2018) should instead be considered a subfamily Baffinelloideae. We also suggest that *Proteomonas* be attributed its own subfamily; Proteomonadoideae.

In Clade 2, *Cryptomonas* (2G) and *Proteomonas* (2B) are reliable genera, and the recently described monospecific genera *Urgorri* and *Baffinella* are sufficiently distinct to justify their independence. Although Clade 2F’s position and origin is uncertain, *Hanusia* and *Guillardia* always group together. However, they are relatively undifferentiated compared to other genera, and perhaps should be merged into a single genus. If so, *Guillardia* Hill and Wetherbee (1990) has priority over *Hanusia* Deane et al. (1998).

Clade 2A contains *Rhodomonas*, *Rhinomonas*, and *Storeatula*, which have been placed together in the family Pyrenomonadaceae (perhaps better termed as a subfamily Rhodomonadoideae), plus a *Teleaulax* and three *Cryptomonas* strains. One possibility is to consolidate members of this clade into a single genus (*Rhodomonas* Karsten, 1898 has priority), but arguments could also be made to divide 2A into two, three, or even four genera. None of these subdivisions map onto taxonomic identifications of the strains in our phylogeny.

Clade 2C also presents challenges. One possibility is merger into a single genus, but it is not clear what that would be. It is clear that the only freshwater strain in this clade, *Rhodomonas minuta* CPCC344, needs to be redescribed. Ideally, the four species of *Teleaulax* that were not included will be in the future, and if so, they may shed light on whether a single genus is warranted, or if *Geminigera* and *Teleaulax* should both be retained, and a new genus defined for the freshwater *R. minuta*.

Several strains do not appear in the phylogeny grouped with their congeners, including *Rhodomonas minuta* (CPCC 344), *Cryptomonas calceiformis* CCAP979/6, and *Teleaulax* sp. (RCC 4857). While *Rhodomonas minuta* (CPCC 344) is the only freshwater *Rhodomonas* strain in this study, it is not the only known Cr-PE545 containing freshwater *Rhodomonas* strain (see for example *Rhodomonas* sp. CCAC 1480B in Marin et al., 1998 and Hoef-Emden, 2008). Furthermore, it is likely that *Rhodomonas minuta* (CPCC 344) is a misidentified *Rhodomonas* based on its grouping in Clade 2C with *Teleaulax* and *Geminigera*. For the marine Cr-PE545-containing *Cryptomonas* strain (*C*. *calceiformis* CCAP979/6) found in Clade 2A along with *Rhodomonas*/*Rhinomonas*/*Storeatula*, this is likely an issue with the early classification scheme developed by Butcher (1967) and followed by Lucas (1968) which uses trichocyst row number and color to delete *Rhodomonas* and divide those marine strains among *Cryptomonas* and *Chroomonas* genera. In fact, *Cryptomonas calceiformis* CCAP979/6 is based on the “Lucas 1968 authority” on the Culture Collection of Algae & Protozoa (CCAP) website. Finally, *Teleaulax* sp (RCC 4857) may be misnamed due to a lack of understanding of the dimorphic life cycle of cryptophytes (see Altenburger et al., 2020).

## Acknowledgements

We would like to thank the three anonymous reviewers for their insightful comments and suggestions. We would also like to thank Jake Swanson and Kristin Heidenreich for isolating novel cryptophyte strains.

**Supplementary Material Fig. S1**. Maximum likelihood phylogeny of nuclear UCE loci. Bootstrap support values are listed for all branches.

**Supplementary Material Fig. S2**. Maximum likelihood phylogeny of nucleomorph UCE loci. Bootstrap support values are listed for all branches. The phylogeny was midpoint rooted.

**Supplementary Material Fig. S3**. Maximum likelihood phylogeny of plastid UCE loci. Bootstrap support values are listed for all branches.

**Supplementary Material Fig. S4**. Multispecies coalescent (MSC) phylogeny. ASTRAL-III local posterior probability are listed for each branch (Zhang et al., 2018). The clade designations match the maximum likelihood phylogeny (Figure 1).

**Supplementary Material Fig. S5**. DiscoVista relative frequency analysis (Sayyari et al., 2018). The phylogeny on the right side is the DiscoVista simplified multispecies coalescent (MSC) phylogeny. For each internal branch of the phylogeny a relative frequency of single-locus trees that support that branch are reported for each single-locus tree dataset (nuclear and nucleomorph + plastid) based on the evolutionary history of cryptophytes. Each internal branch has four connected branches that form three different topologies. The red (t1) bars indicate relative frequency of single-locus trees that support the maximum likelihood phylogeny, and the blue (t2 and t3) bars are alternative topologies. The dotted line indicates one-third of the total number of single-locus trees. Each topology is detailed on the x-axis where the branches separated by a comma are joined.

**Supplementary Table S1**. List of cryptophyte strains used in this study. Data includes culture center or field collection site, growth media, growth temperature, salinity (freshwater or marine), cryptophyte phycobiliprotein (PBP) type, and PBP maximum wavelength (if available). Strains with cryptophyte PBP data from Cunningham et al. (2019) are in bold text and PBP data for *Hemiselmis aquamarina* (RCC 4102) is from Magalhaes et al. (2021).

**Supplementary Table S2**. Results of nucleotide substitution saturation assessment produced using PhyloMAD for each UCE locus (Duchene et al., 2018).

**Supplementary Table S3**. List of cryptophyte genera missing from this study. Data includes number of known species isolates, Web of Science (WOS) records, AlgaeBase (AB) records, habitat (freshwater or marine), family, holotype, and additional notes.

