## Supplementary figures and images for "A three-genome ultraconserved element phylogeny of Cryptophytes"

### Supplementary Figure S1

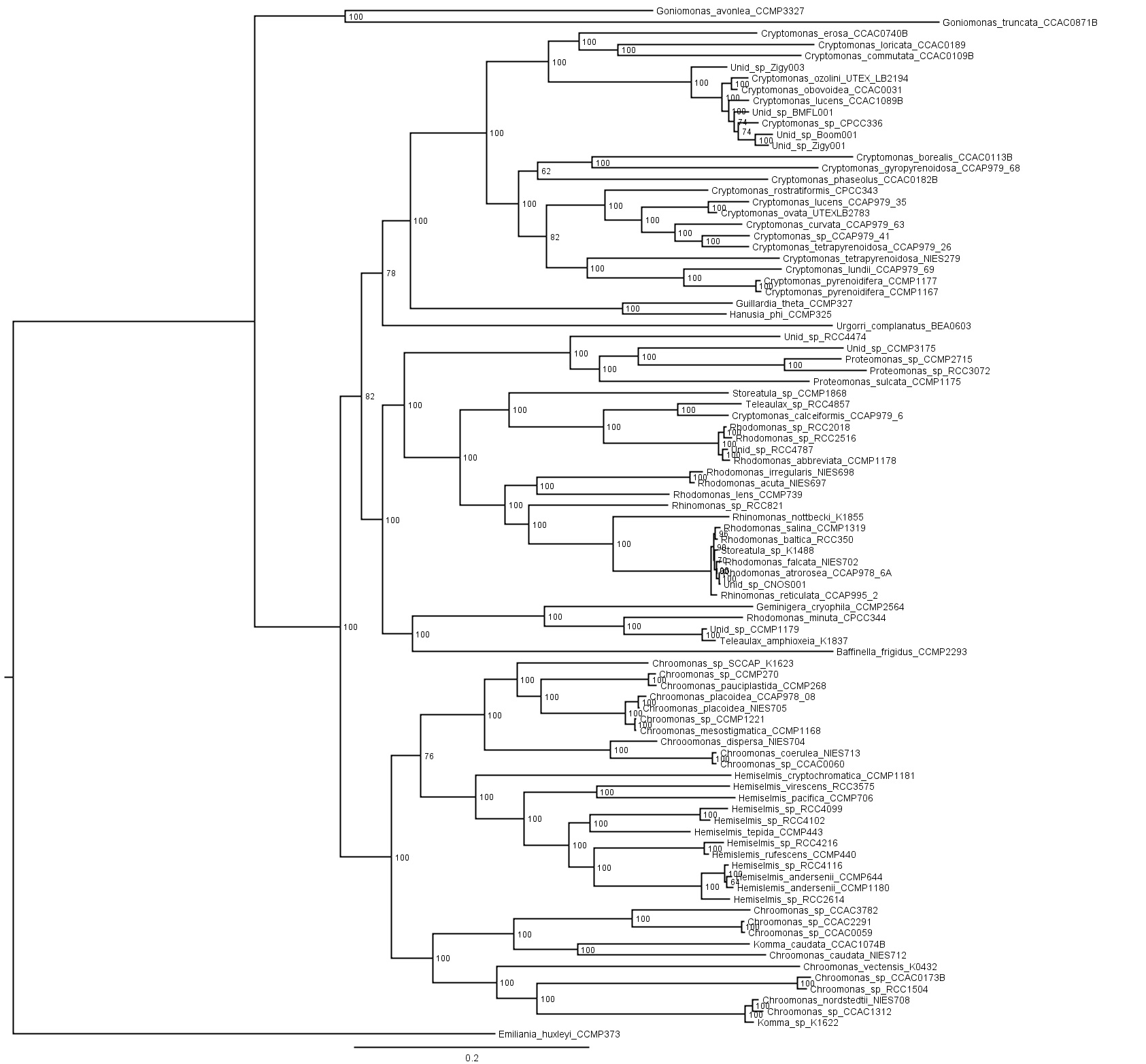

### Supplementary Figure S2

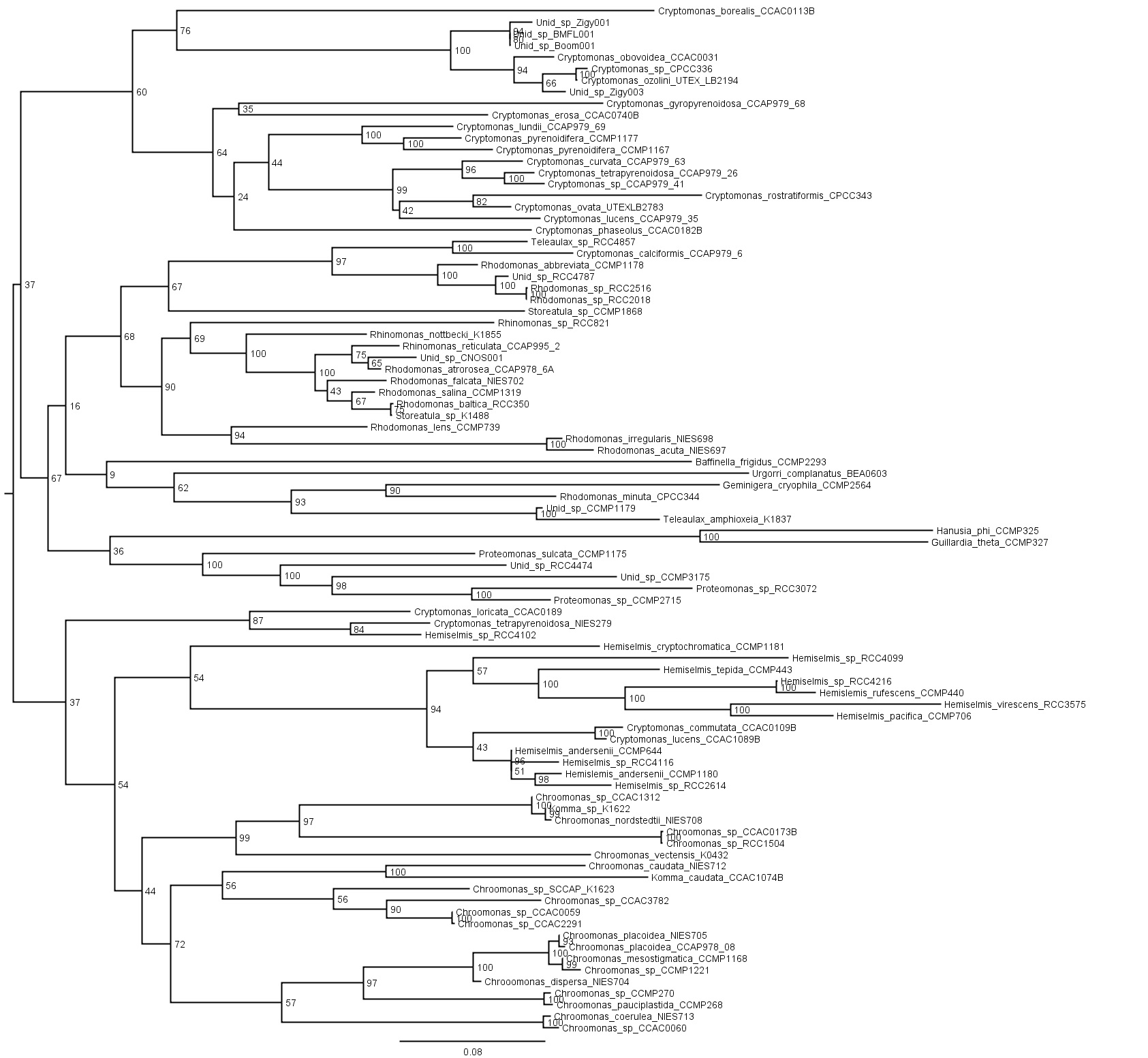

### Supplementary Figure S3

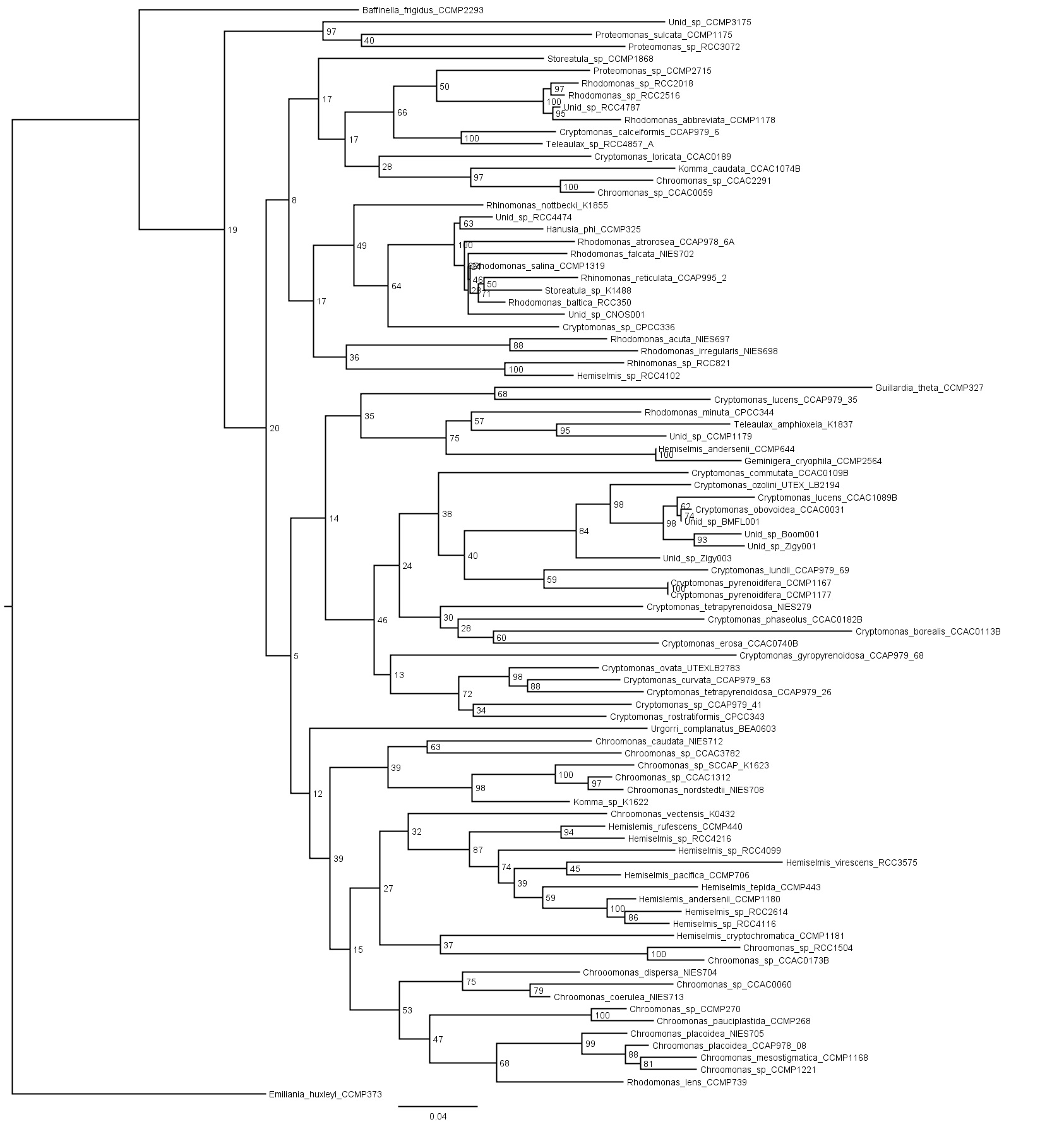

### Supplementary Figure S4

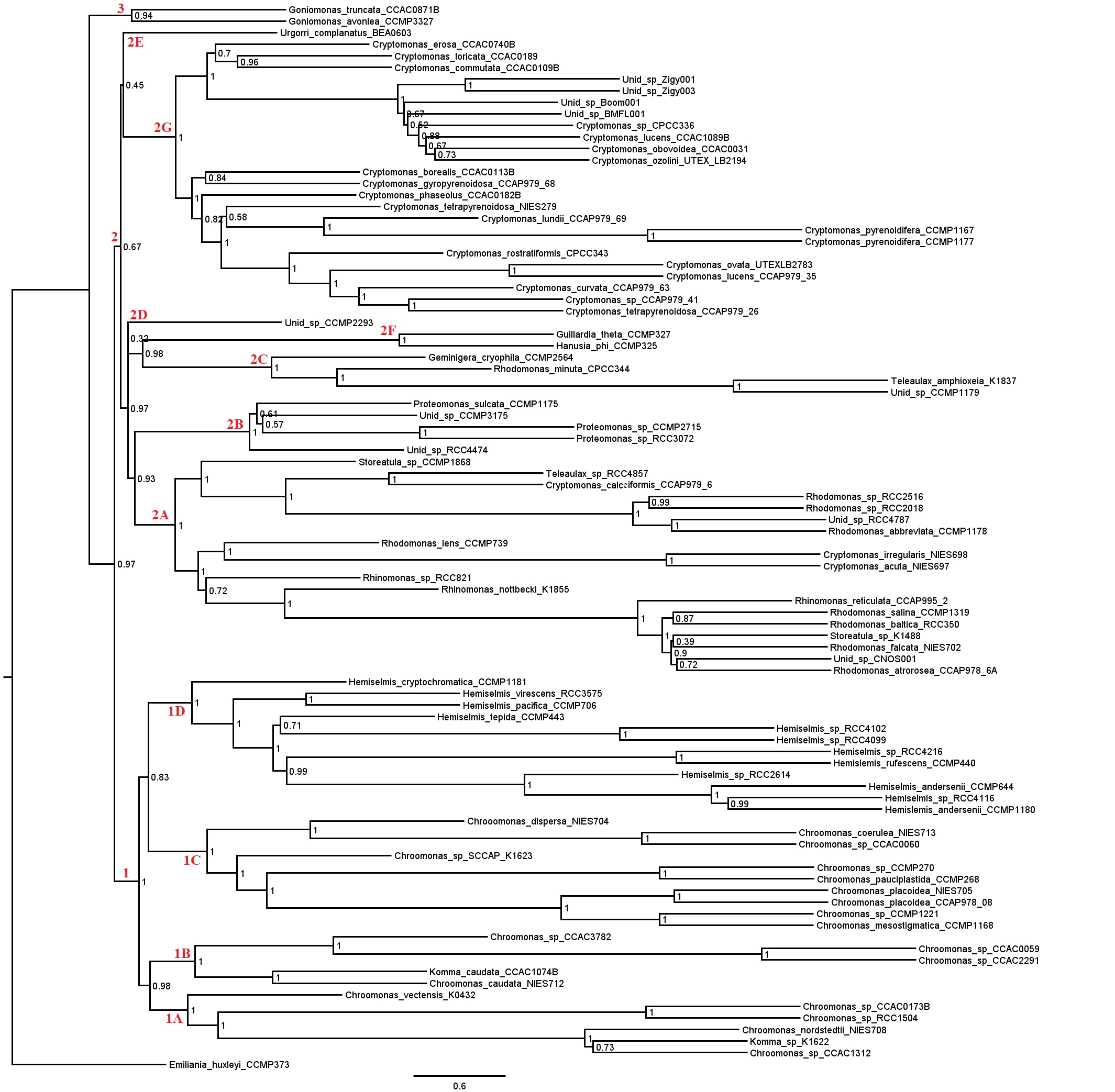

### Supplementary Figure S5

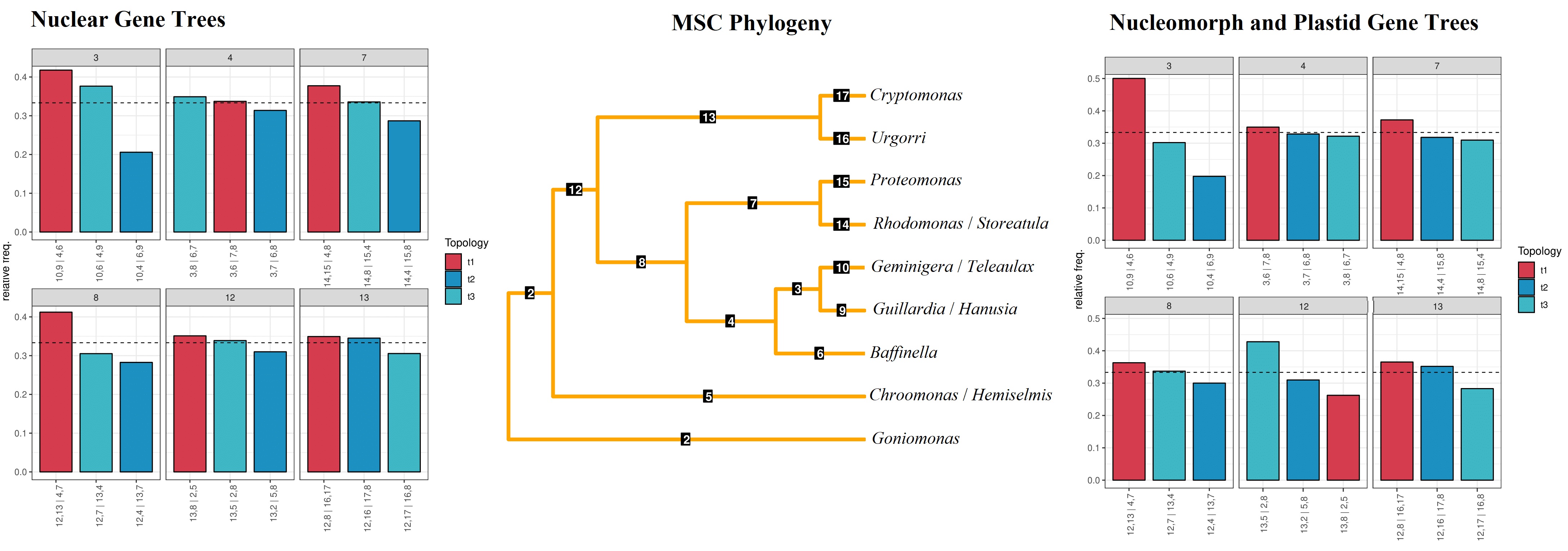
